# Neuroinflammation in post-acute sequelae of COVID-19 (PASC) as assessed by [^11^C]PBR28 PET correlates with vascular disease measures

**DOI:** 10.1101/2023.10.19.563117

**Authors:** Michael B. VanElzakker, Hannah F. Bues, Ludovica Brusaferri, Minhae Kim, Deena Saadi, Eva-Maria Ratai, Darin D. Dougherty, Marco L. Loggia

**Affiliations:** Division of Neurotherapeutics, Department of Psychiatry, Athinoula A. Martinos Center for Biomedical Imaging, Massachusetts General Hospital, Harvard Medical School, Boston, MA, USA; Department of Radiology, Athinoula A. Martinos Center for Biomedical Imaging, Massachusetts General Hospital, Harvard Medical School, Boston, MA, USA; Department of Computer Science And Informatics, School of Engineering, London South Bank University, London, UK; Department of Anesthesia, Critical Care and Pain Medicine, Massachusetts General Hospital, Harvard Medical School, Boston, MA, USA; PolyBio Research Foundation, Medford, MA, USA

**Author notes:** Correspondence to: Michael VanElzakker PhD, 149 Thirteenth Street, Office 2610, Building 149, Martinos Center for Biomedical Imaging Charlestown MA 02129.

**Keywords:** COVID-19, Long COVID pathogenesis, neuroimaging, positron emission tomography, fibrinogen, cardiovascular, glia, microglia, brain inflammation

## Abstract

The COVID-19 pandemic caused by SARS-CoV-2 has triggered a consequential public health crisis of post-acute sequelae of COVID-19 (PASC), sometimes referred to as long COVID. The mechanisms of the heterogeneous persistent symptoms and signs that comprise PASC are under investigation, and several studies have pointed to the central nervous and vascular systems as being potential sites of dysfunction. In the current study, we recruited individuals with PASC with diverse symptoms, and examined the relationship between neuroinflammation and circulating markers of vascular dysfunction. We used [^11^C]PBR28 PET neuroimaging, a marker of neuroinflammation, to compare 12 PASC individuals versus 43 normative healthy controls. We found significantly increased neuroinflammation in PASC versus controls across a wide swath of brain regions including midcingulate and anterior cingulate cortex, corpus callosum, thalamus, basal ganglia, and at the boundaries of ventricles. We also collected and analyzed peripheral blood plasma from the PASC individuals and found significant positive correlations between neuroinflammation and several circulating analytes related to vascular dysfunction. These results suggest that an interaction between neuroinflammation and vascular health may contribute to common symptoms of PASC.

## 1. INTRODUCTION

The coronavirus disease 2019 (COVID-19) global pandemic caused by severe acute respiratory syndrome coronavirus 2 (SARS-CoV-2) has triggered a consequential public health crisis of long COVID or post-acute sequelae of COVID-19 (PASC), defined by symptoms that begin following acute SARS-CoV-2 infection or persist from that initial acute illness (Centers for Disease Control and Prevention, 2022; World Health Organization, 2021).

The US Census Bureau’s Household Pulse Survey found that, as of June 2023, about 11% of adults who had ever had COVID-19 reported that they were currently experiencing symptoms of PASC or long COVID, with nearly 16% of adults stating that they had PASC or long COVID symptoms at some point in time (National Center for Health Statistics, 2023). PASC is an umbrella term applied to a heterogeneous group of patients. Symptoms can be diverse and commonly include nonspecific neurological and neuropsychiatric symptoms including difficulty concentrating or subjective ‘brain fog,’ unusually profound fatigue, headaches, depression, anxiety, body pain, and disrupted sleep (Davis et al., 2021; Zeng et al., 2023). These symptoms can significantly impact quality of life (Líška et al., 2022), and can be initiated even by mild acute COVID-19 illness (Boribong et al., 2022; Ma et al., 2023).

PASC mechanisms remain under study, however SARS-CoV-2 is now understood to infect a wide range of tissue types and is capable of driving inflammation, coagulation, and vascular problems (Klein et al., 2022; Pretorius et al., 2022; Proal & VanElzakker, 2021). Early evidence demonstrates that vascular related problems seen in acute COVID-19 illness (Varga et al., 2020) may persist in some individuals (Xie et al., 2022), and thus may contribute to PASC symptoms. The current study investigated whether PASC patients with diverse symptom presentation showed increases in a measure of neuroinflammation relative to healthy controls with no known history of COVID-19, and whether PASC neuroinflammation was related to measures of vascular health.

As a densely vascularized organ that uses 15-20% of the body’s total circulating blood supply (Berkman et al., 2021), the brain is uniquely vulnerable to disruptions to vascular health. COVID-19 appears to confer a potent vulnerability to neurovascular problems. For example, in patients that survived acute COVID-19, in the following year the risk for hemorrhagic stroke and for cerebral venous thrombosis more than doubled (Xie et al., 2022; Xu et al., 2022). As evidence of general consequences for the brain, a longitudinal study comparing pre-pandemic structural neuroimaging data to post-COVID structural neuroimages from the same individual showed small but significant reduction of grey matter thickness and whole-brain volume as well as increased markers of tissue damage, relative to longitudinal scans from individuals not infected with SARS-CoV-2 (Douaud et al., 2022). In PASC, a neuroimaging study using arterial spin labeling fMRI found evidence of decreased neurovascular perfusion in patients reporting persistent cognitive problems (Ajčević et al., 2023), potentially relating central nervous system and vascular dysfunctions. Despite these observations, no PASC study has yet directly linked neuroinflammation and vascular dysfunction.

In addition to their role as the innate immune cells of the central nervous system, glial cells are critical to normal central nervous system functioning, including playing a central role in neurotransmission, neurovascular function, and blood-brain barrier integrity. Thus, the activity of glial cells may be an important connection between brain and vascular abnormalities in PASC (Cabezas et al., 2014; Mestre et al., 2017). Like other innate immune cells, glia can enter a spectrum of ‘activated’ states when they detect paracrine inflammatory mediators (e.g. proinflammatory cytokines), pathogen- associated molecular patterns (PAMPs), or damage-associated molecular patterns (DAMPs). Short-term glial activation is central to their role as cells of innate, or nonspecific, immunity that clear multiple forms of pathogens or cell debris. The glial role in the innate immune system also involves general ‘sickness’ symptoms that shift an organism’s behavior to preserve energy at a time of high energy demand (Dantzer & Kelley, 2007; VanElzakker et al., 2019). However, long-term glial activation can disrupt the delicate symbiosis of glia, blood vessels, neurons, and cerebrospinal fluid in normal central nervous system function, and ongoing glial activation occurs in multiple neurological and neuropsychiatric conditions (VanElzakker et al., 2019) and is associated with symptoms that are commonly reported in PASC such as disrupted cognition (Barrientos et al., 2006; Lindgren et al., 2020), increased pain (Loggia et al., 2015), and other nonspecific symptoms of sickness.

Activation of glia is a key component of neuroinflammation, which can include other mechanisms such as activated peripheral immune cells penetrating into brain, inflammatory activation of neurovascular endothelial cells, or density of motile microglia. Rodent models have shown upregulation of the translocator protein (TSPO) during glial activation states (Pannell et al., 2020), and PET neuroimaging with TSPO-specific radioligands is a widely used method in the study of human neuroinflammation (Albrecht et al., 2016; VanElzakker et al., 2019). [^11^C]PBR28 is a second-generation TSPO-binding radioligand that has good specificity for TSPO and a low background signal-to-noise ratio (Albrecht et al., 2016). Thus, the measurement of central nervous system TSPO concentration in concert with analysis of circulating measures related to vascular damage may provide insight into PASC mechanisms.

## 2. MATERIALS AND METHODS

### 2.1 STUDY DESIGN

In this case-control, cross-sectional study, we compared 12 PASC versus an existing dataset of 43 control individuals with no known prior COVID-19 exposure. We used PET neuroimaging to study neuroinflammation (here operationalized as increased TSPO radioligand binding) in PASC by comparing [^11^C]PBR28 standardized uptake value ratios (SUVR) in PASC versus controls. All participants answered questionnaires related to pain and depression; PASC participants answered additional questionnaires related to their PASC symptoms and history.

We also analyzed peripheral blood collected from PASC participants immediately before their PET scan to study the relationship between central nervous system glial activation in PASC and measures related to vascular health, inflammation, and angiogenesis. For the blood measurements, platelet-poor plasma (PPP) was collected from 11 PASC cases immediately before [^11^C]PBR28 injection.

### 2.2 PARTICIPANTS

As PASC is an umbrella term with a heterogeneous presentation, cohort phenotyping is key to study design. To prioritize recruitment of individuals with diverse nonspecific symptoms, we used a modified myalgic encephalomyelitis/chronic fatigue syndrome (ME/CFS) International Consensus Criteria (ICC, Carruthers et al., 2011) as PASC group inclusion criteria, requiring at least one symptom from each of the 4 ICC symptom clusters: post-exertion exhaustion, neurological impairments, immune/gastro- intestinal/genito-urinary, and energy/autonomic. All PASC participants had onsetting COVID-19 illness that occurred before August 2021 and at least 10 months prior to the scan date (mean=20.50 months, SD=7.75), reflecting pre-Omicron strains and qualifying them for both CDC and WHO definitions of PASC (4 weeks and 3 months of symptoms, respectively). Two of the 12 PASC participants were hospitalized during their acute COVID-19 (without being put on a ventilator), and 1 of 12 reported being vaccinated prior to the acute COVID-19 infection that initiated their PASC.

A total of 12 PASC study participants were compared to an existing normative dataset of 43 healthy controls with no known history of COVID-19, that were scanned with the same protocol in the same scanner, between 2013-2021. Control scans were selected from multiple completed or ongoing studies (e.g., Albrecht et al., 2019; Alshelh et al., 2020; Loggia et al., 2015); 34 of the 43 controls were scanned pre-pandemic and all post-pandemic-onset controls had a negative plasma antibody test but were scanned in Massachusetts’ state-of-emergency period of the pandemic (Brusaferri et al., 2022). Participants of any sex were recruited to participate in a ∼90 minute scanning session at the HST/MGH (Harvard-Massachusetts Institute of Technology Program in Health Sciences and Technology/Massachusetts General Hospital) A.A. Martinos Center for Biomedical Imaging, in Boston MA.

Exclusionary criteria included PET or MRI contraindications (e.g., metallic implants, surpassed FDA research-related PET scan limit in past 12 months, major kidney or liver problems, pregnancy), history of other neurological disorders (epilepsy, history of stroke, tumor, brain tissue-damaging pathologies), history of major head trauma (loss of consciousness for more than 5 minutes), type 1 diabetes, major cardiac event in the past decade, current acute illness or infection (e.g., COVID-19, cold, flu), or history of psychotic disorder or other major psychiatric illness except posttraumatic stress disorder (PTSD), depression, and anxiety which were exclusionary only if the conditions were so severe as to require hospitalization in the past 5 years. Before the scan day, patients were genotyped via saliva or blood for the Ala147Thr polymorphism in the *TSPO* gene (rs6971 polymorphism), which is known to affect binding affinity for several TSPO radioligands, including [^11^C]PBR28 (Owen et al., 2012, 2015). Individuals with the Ala/Ala or Ala/Thr genotypes (predicted high- and mixed-affinity binders, respectively) were included, and the genotype was modeled as a covariate in the statistical design. Individuals with the Thr/Thr genotype (predicted low-affinity binders) were excluded at the time of the screening and therefore not represented in our dataset. Medication exclusions included use of immunosuppressive medications such as prednisone or TNF blocking medications within the 2 weeks preceding the visit, routine use of benzodiazepines or any use in the past 2 weeks except clonazepam (Klonopin), lorazepam (Ativan), and alprazolam (Xanax), which have documented or predicted low affinity for TSPO (e.g., Kalk et al., 2013).

Before the scan, all participants underwent a history and physical examination by an experienced nurse practitioner to confirm PET-MRI safety. Participants answered several questionnaires and their height and weight were recorded. Immediately before neuroimaging, IV blood was drawn. Blood-based pregnancy testing (hCG) was performed on all individuals capable of becoming pregnant.

### 2.3 PET IMAGING PARAMETERS

Our dual PET-MRI scanner consists of a 3-Tesla (3T) Siemens TIM Trio MRI (60 cm bore diameter) with 8-channel head coil and a photodiode-based high-resolution BrainPET head camera insert. The BrainPET prototype is a dedicated brain scanner that has 32 detector cassettes, each consisting of 6 detector blocks, each made up of a 1212 array of lutetium oxyorthosilicate crystals (2.52 mm^3^) read out by magnetic field- insensitive avalanche photodiodes. The transaxial and axial fields of view are 32 cm and 19.125 cm, respectively. The 3T MR system is equipped with the standard “TIM” (total imaging matrix) 32 RF channel receivers, accommodating up to 32 element array coils. [^11^C]PBR28 radioligand was produced by an in-house Siemens Eclipse HP self-shielded 11 MeV cyclotron with single-beam extraction and a four position target- changer. During the PET scan, [^11^C]PBR28 (up to 15 mCi, which is equivalent to ∼3.7 mSv) was injected intravenously with a slow bolus over a ∼30 second period (Debruyne et al., 2003). The catheter was then flushed post-injection with 0.9% sterile saline solution. Dynamic data was collected by the head PET camera over 90 minutes list mode and framed post-collection; concurrent high-resolution T1-weighted (MEMPRAGE) structural MRI images were collected for spatial registration of the PET data, as well as generation of attenuation correction maps (Izquierdo-Garcia et al., 2014) and scatter correction.

### 2.4 BLOOD COLLECTION

Immediately before [^11^C]PBR28 injection and scanning, venous blood was collected from 11 PASC participants into a citrated vacutainer tube (BD 369714 light blue cap) (one participant was scanned when a study team member had COVID-19 and therefore blood was not collected). The citrate tube was gently upended 3x then left to sit at room temperature for 30 minutes, before centrifugation at 15min x 3000 RCF x 20°C. Platelet-poor plasma (PPP) supernatant was pipetted into sterile 1.5mL microtubes (Sarstedt Biosphere 72.703.217) and stored at -80°C. For analysis, these samples from the 11 PASC participants were thawed on wet ice and 275µL aliquots were pipetted into sterile SureSealTM 0.65mL microfuge tubes (VWR Ward’s Science 470228-444), marked with de-identified alphanumeric sequences, and sent overnight on dry ice to Eve Technologies (Calgary Canada) for analysis.

Three multiplex panels were conducted based upon previous PASC literature: vascular health, cytokine, and angiogenesis (e.g., Proal & VanElzakker, 2021); Klein et al., 2022; Patel et al., 2022). Analytes per multiplex were as follows: Vascular health (α2- macroglobulin, orosomucoid, CRP [C-reactive protein], fetuin A, fibrinogen, haptoglobin, sL-selectin, PF4 [platelet factor 4], pentraxin-2) (Millipore HCVD3MAG-67K Luminex magnetic bead panel); Cytokines (GM-CSF, IFNγ, IL-1β, IL-1RA, IL-2, IL-4, IL-5, IL-6, IL-8, IL-10, IL-12(p40), IL-12(p70), IL-13, MCP-1, TNFα) (Millipore HCYTA-60K Luminex magnetic bead panel); Angiogenesis (angiopoietin-2, BMP-9, EGF, endoglin, endothelin- 1, FGF-1, FGF-2, follistatin, G-CSF, HB-EGF, HGF, IL-8, leptin, PLGF, VEGF-A, VEGF-C, VEGF-D) (Millipore HAGP1MAG-12K Luminex magnetic bead panel). Concentrations of these analytes were entered into a correlation matrix to test their relationship to extracted SUVR values and symptom measures in the PASC participants. Items from the vascular health panel are described in Box 1.

### 2.5 BEHAVIORAL MEASURES

All 12 PASC and 43 eligible control participants were administered items from the Brief Pain Inventory (BPI) (Cleeland & Ryan, 1994) and the Beck Depression Inventory (BDI) (Beck et al., 1996).

PASC participants were screened to fulfill a modified myalgic encephalomyelitis/chronic fatigue syndrome (ME/CFS) International Consensus Criteria (ICC) criteria, and then on the day of the scan were asked to rate each ICC symptom’s severity from 0-10. PASC individuals were also given a revised medical history questionnaire that asked about the history of several neurological or neuropsychiatric symptoms by naming a symptom then giving the option to either leave blank or select one of three options: 1) This was a problem for me before COVID-19, 2) This is a problem for me since I had COVID-19, or 3) This is a serious problem for me since I had COVID-19.

### 2.6 PET DATA PROCESSING

For the PASC versus control PET comparison, PET data processing was performed on a custom Linux-based pipeline used in several previous studies (e.g., Albrecht et al., 2019; Alshelh et al., 2021; Brusaferri et al., 2022; Loggia et al., 2015), including standard preprocessing and PET-MRI spatial registration to MNI (Montreal Neurological Institute) space.

Following data acquisition, several quality control steps were performed including by-participant visual inspection of detector blocks (Tayal et al., 2021), spatial registration, skull stripping, artifacts, and attenuation image map alignment. After normalizing radioactivity by injected dose by body weight, PET data were scatter- and attenuation-corrected using a custom MR-based method developed at the Martinos Center for Biomedical Imaging (Izquierdo-Garcia et al., 2014). For all subjects, 30-minute static standardized uptake value (SUV) PET images were reconstructed from data acquired ∼60-90 min after the injection of the tracer. SUV maps were spatially normalized by nonlinear transformation into common MNI space and smoothed with an 8mm FWHM (full-width half-maximum) Gaussian kernel. Standard uptake value ratio (SUVR) images were then created by intensity-normalizing individual MNI-space SUV maps to cerebellum uptake, anatomically-defined by AAL (automatic anatomical labeling) atlas (Tzourio-Mazoyer et al., 2002). There were no group differences in cerebellum SUV in the 12 vs. 43 primary or the 11 vs. 11 validation analyses (t(53)=0.82,p=.42; t(10)=0.85,p=.41 respectively).

### 2.7 STATISTICAL ANALYSIS

We conducted a primary unpaired (between groups) comparison including all available PET data (12 PASC vs. 43 control). Cerebellum-normalized, MNI-registered SUVR maps for each participant were entered into a general linear model, using the FEAT software tool (version 6.00, Woolrich et al., 2004) within FSL (FMRIB Software Library, Wellcome Centre). Nonparametric permutation inference of [^11^C]PBR28 SUVR images was performed using FSL Randomise, with 5000 permutations (Winkler et al., 2014) and statistically controlling for sex and TSPO polymorphism. Cluster-correction was used to account for multiple comparisons, using a cluster-forming threshold of Z=2.3 and cluster size of p=.05 . Demographic and PET-relevant data comparisons for the primary analysis are shown in Table 1. Reflecting the sex distribution of PASC (Bai et al., 2022), there was a relatively high proportion of females (10) in the PASC group.

**Table 1.** In the primary PET comparison (i.e. n=12 PASC vs. n=43 control), an unpaired t-test or Chi-square revealed no significant difference between PASC and control groups in age, genotype frequency, BMI, or injected dose. Reflecting the sex distribution of PASC (Bai et al., 2022), there was a higher proportion of female participants in the PASC group relative to the larger historical control group. In this primary analysis, we statistically controlled for genotype and sex.

|  | PASC | Control | Statistics |
| --- | --- | --- | --- |
| Age mean (SD) | 47.25 yr (14.19) | 50.86 yr (14.19) | $t(53)=0.84$, $p=.20$ |
| Sex | F=10, M=2 | F=17, M=26 | $\chi^2(1.55)=7.20$, $p=.007$ |
| Genotype count | GG=5, GA=7 | GG=27, GA=16 | $\chi^2(1.55)=1.72$, $p=.19$ |
| BMI mean (SD) | 27.25 (5.01) | 26.19 (4.96) | $t(53)=0.66$, $p=.51$ |
| Injected Dose | 13.64 mCi (1.90) | 13.26 mCi (1.58) | $t(53)=0.70$, $p=.49$ |

While we did statistically control for sex in the primary analysis, we performed a second validation analysis using a paired approach in which each PASC participant was matched to a control participant for genotype, sex, and age ±5 years (Supplemental data).

To test the relationship between SUVR and blood analytes or questionnaire data, simple Pearson’s correlations were calculated. For these correlation analyses, we used the significant cluster that arose from the primary 12 PASC vs 43 control comparison (see Results) and extracted each participant’s average SUVR value from that cluster. For behavioral/demographic data, group statistical comparisons were performed using unpaired or paired Student’s t-tests, Chi-square tests, or Pearson’s correlation as appropriate, while BDI subscales were analyzed with a mixed-model ANOVA. Two-tailed p-values are reported in all cases. Cohen’s D effect size for the primary analysis was calculated using each participant’s mean SUVR from the whole-brain significant cluster.

## 3. RESULTS

### 3.1 IMAGING RESULTS: VOXEL-WISE GROUP DIFFERENCES

The voxelwise whole-brain comparison of PASC cases versus healthy controls revealed a significant increase in [^11^C]PBR28 PET signal at the Z=2.3 threshold (large Cohen’s D effect size of 1.28). This increased PET signal spanned a wide swath of brain regions including midcingulate cortex, corpus callosum, thalamus, basal ganglia/striatum, subfornical organ, anterior cingulate cortex, medial frontal gyrus, and precentral gyrus (see Figure 1 and Table 2), and is suggestive of glial activation or neuroinflammation.

**Figure 1.**
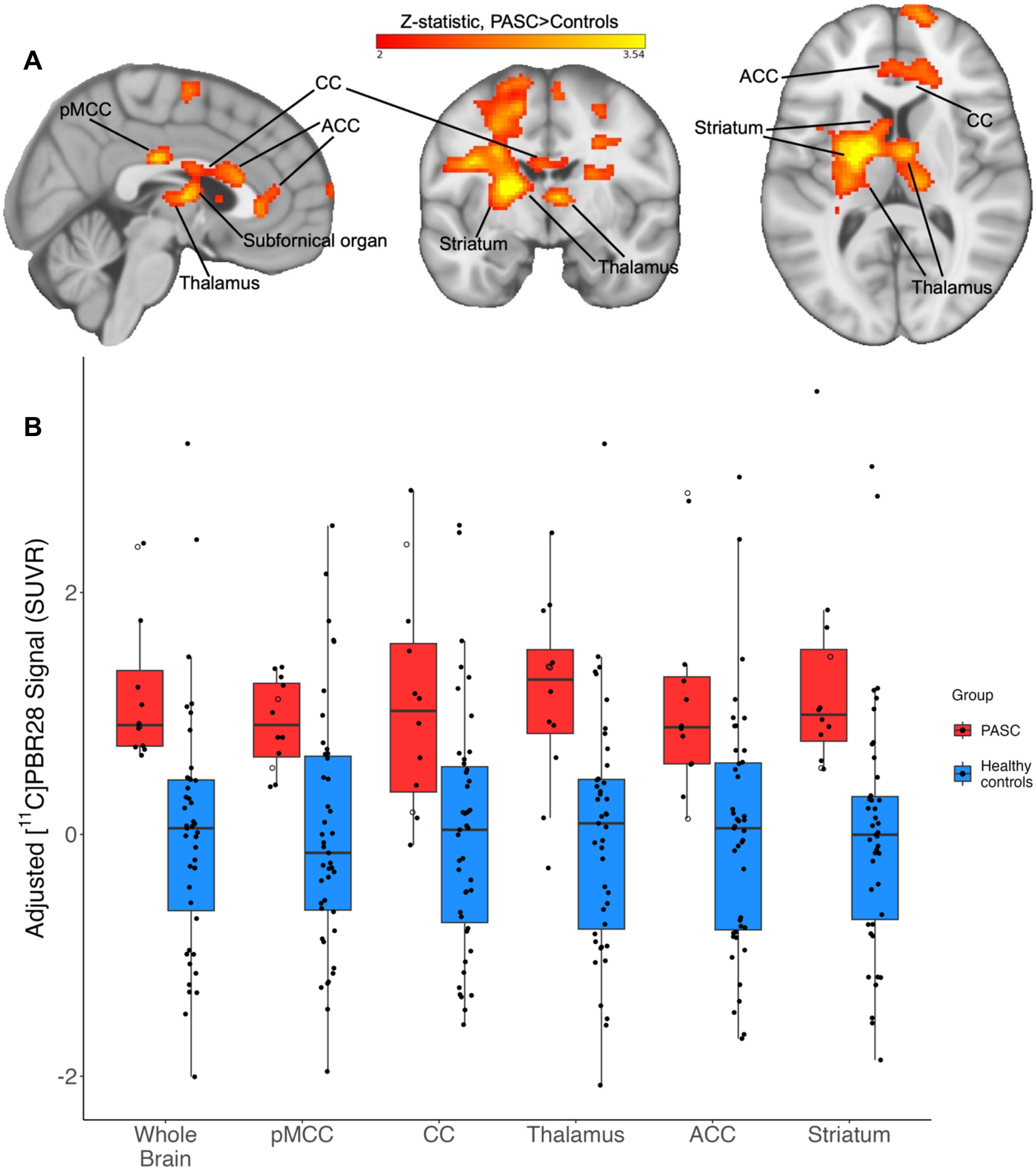
A: Example slices through sagittal (x=3), coronal (y=-7), and axial (z=10) sections of the 12 PASC > 43 control group-level unpaired (between-groups) comparison, showing the pattern of increased [^11^C]PBR28 signal (shown in neurological convention). Color bar: threshold min. Z score of 2 and max. 3.54. B: PET signal data extracted from individual study participants, depicted from the primary whole-brain analysis and across five example bilateral brain regions. Y-axis data points represent the mean SUVR values of significant voxels within each significant cluster region, with each structure’s data normalized to the control average. PASC participants hospitalized during their acute COVID-19 illness are indicated with an open circle. Brain regions are anatomically defined by the Harvard-Oxford Atlas (thalamus, ACC, striatum) or CC by the Jüelich Atlas at >70% probability threshold. In the case of the pMCC, we used the Harvard-Oxford posterior cingulate anatomical mask, but the activation cluster was only within pMCC. In the PASC group, the two hospitalized cases are depicted by open circles. pMCC = posterior midcingulate cortex; CC = corpus callosum; ACC = anterior cingulate cortex

**Table 2.** : Peak voxel coordinates from brain regions significantly increased in PASC versus controls in whole-brain [^11^C]PBR28 voxelwise comparison.

| Region with local peak | Z value | MNI coordinates (x,y,z) |
| --- | --- | --- |
| Posterior midcingulate | 3.54 | 4, -22, 28 |
| Corpus callosum | 2.94 | 3, 20, 18 |
| Thalamus (L) | 3.54 | -20, -2, 8 |
| Basal ganglia (L) | 3.54 | -14, -6, 15 |
| Superior frontal gyrus (R) | 3.54 | 14, 38, 40 |
| Subfornical organ (estimated) | 3.09 | 4, -7, 8 |
| Anterior midcingulate | 2.95 | 4, 19, 19 |
| Precentral gyrus (R) | 3.54 | 36, 1, 36 |
| Postcentral gyrus (R) | 3.54 | 40, -18, 40 |
| Retrolenticular internal capsule (L) | 3.54 | -28, -38, 18 |
| Superior corona radiata (L) | 3.54 | -26, -6, 32 |

There were no voxels with significantly greater SUVR values in the controls > PASC contrast.

In order to visualize data distribution, individual participants’ data were extracted from the primary voxelwise analysis’ significant cluster, and from a selection of atlas-defined anatomical regions that showed significant signal. The distribution pattern of extracted SUVR values showed that this effect was not driven by outliers or by hospitalized cases (Figure 1B).

We conducted a paired validation analysis with individually matched PASC-control pairs and found a very similar PET signal pattern (Supplemental Figure 1). In this analysis the average injected radioligand dose also did not differ between PASC (mean 13.85 mCi, SD 1.83) and controls (mean 12.63 mCi, SD 1.47), t(10)=1.69, p=.12.

### 3.2 BLOOD ANALYTE MEASURES and CORRELATIONS

We found positive Pearson’s r correlations between PET signal and the majority of analytes from the vascular disease multiplex panel (see Figure 2, Table 3). Specifically, we found positive moderate to strong correlations between the PET signal extracted for each individual from the cluster derived from the voxelwise group comparison, and the concentrations of seven plasma vascular health-associated factors: fibrinogen (r(9)=.80, p=.0032), α2-macroglobulin (r(9)=.73, p=.011), orosomucoid (alpha-1-acid glycoprotein or AGP) (r(9)=.69, p=.019), fetuin A (r(9)=.68, p=.022), sL-selectin (soluble leukocyte selectin, or sCD62L) (r(8)=.70, p=.025), pentraxin-2 (serum amyloid P component, or SAP) (r(9)=.66, p=.026), and haptoglobin (r(8)=.74, p=.015). The haptoglobin scatterplot appeared to include a potential outlier. However, removing this participant from the analyses did not substantially affect significance, (r(7)=.88, p=.0018). The correlation with the C-reactive protein (CRP) was trending but not statistically significant (r(8)=0.61, p=.064). In this case, the relevant scatterplot appeared to include one outlier, as well as another participant’s levels that were not detected by the assay. The CRP correlation remained non-significant after removing the potential outlier from the CRP analysis (p>.4). The high-end outlier in both haptoglobin and CRP analyses was the same participant, who was hospitalized during their acute COVID-19 illness. For both the sL-selectin and haptoglobin assays the same participant’s levels were not detected by the respective assay, and platelet-factor 4 (PF4) was not detected by the assay in the majority of samples.

**Figure 2:**
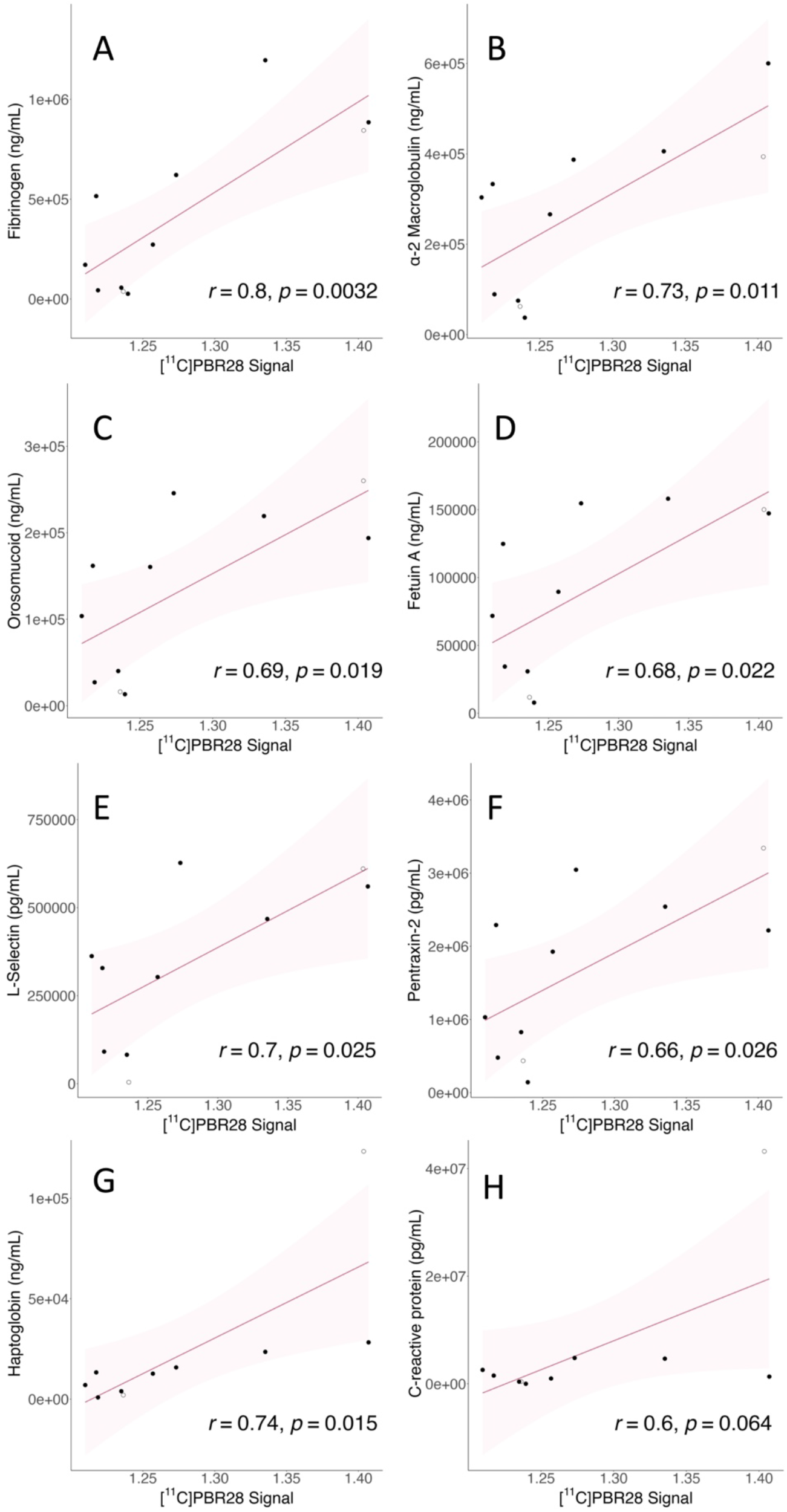
Correlations between vascular health analytes and PET signal. Y-axis shows analyte concentration; units vary. X-axis shows mean PET SUVR values extracted from the whole-brain significant cluster of each individual. Shadow represents the 95% confidence interval, all p-values are reported two-tailed. See Box 1 for analyte descriptions and Table 2 for Pearson’s r values. a. Fibrinogen; b. α-2 macroglobulin; c. Orosomucoid; d. Fetuin A; e. sL-selectin; f. Pentraxin-2; g. Haptoglobin; h. C-reactive protein

**Table 3:** Vascular disease-related blood analytes that significantly correlated with whole-brain [^11^C]PBR28 uptake values extracted from the significant cluster in the PASC group.

| Peripheral blood vascular related analyte | Whole brain PET signal correlation |
| --- | --- |
| Fibrinogen | r(9)=.80, p=.0032 |
| α2-macroglobulin | r(9)=.73, p=.011 |
| Orosomucoid (alpha-1-acid glycoprotein or AGP) | r(9)=.69, p=.019 |
| Fetuin A | r(9)=.68, p=.022 |
| sL-selectin (soluble leukocyte selectin) | r(8)=.70, p=.025 |
| Pentraxin-2 | r(9)=.66, p=.026 |
| Haptoglobin | r(8)=.74, p=.015 |

Cytokine concentrations from the 15-plex panel frequently correlated with one another but did not correlate with PET signal (Supplemental Figure 2), which was our a-priori hypothesis (VanElzakker et al., 2019). Analytes from the angiogenesis 17-plex panel analytes also did not correlate with PET signal. We did not have strong a-priori hypotheses regarding correlations between blood analytes and self-reported symptoms.

### 3.3 SYMPTOM SEVERITY AND HISTORY

Response scores to the BPI severity item “Pain at its worst in the past 24 hrs” were higher in PASC (M=7.08, SD=3.72) versus controls (M=0.44, SD=0.81), t(52)=10.91, p<.001, two-tailed, unequal variance. Mean scores from BPI-Interference subscale items were higher in PASC (M=21.75, SD=15.42) versus controls (M=0.90, SD=0.31), t(51)=9.21, p<.001, two-tailed, unequal variance.

Mean total BDI scores were higher in cases (M=11.58, SD=1.74) versus controls (M=1.14, SD=5.58), t(52)=9.96, p<.001, two-tailed, unequal variance. Mean depression severity according to the BDI was on average categorized as mild (10-18) in PASC and none-to-minimal (0-9) in controls. Two PASC cases met the threshold of moderate depression (i.e., 19-29), with scores of 19 and 25. Of the 12 PASC cases, six reported that depression was not a problem, five reported “This is a problem for me since I had COVID-19” and one reported “This is a serious problem for me since I had COVID-19” about depression.

BDI responses can be divided into affective (mood-centered) versus somatic (body-centered) subscales. To better understand which questionnaire items were driving the higher BDI scores in PASC vs. controls, we conducted a 2x2 mixed model ANOVA with diagnosis (PASC, control) as the between-groups factor and BDI subscale (affective, somatic) as the within-subjects factor. This ANOVA revealed a significant interaction between diagnosis and BDI subscale F(1,52)=51.734, p<.001, (partial eta^2^=.499, large effect size) driven by higher average item somatic subscale scores within the PASC group (Supplemental Table 1). There were also main effects of diagnosis F(1,52)=155.184, p<.001 (consistent with the t-test) as well as a main effect of BDI subscale F(1,52)=54.236, p<.001 in which average item score was higher in somatic versus affective subscales independent of diagnosis.

On the day of the scan, PASC participants self-reported the severity of each of the 21 ICC symptoms on a 0-10 scale (see Supplemental Figure 3); all reported 5 out of 10 severity or greater on at least two of the eight ICC Neurological symptoms cluster items: headache, unrefreshing sleep, significant pain, short-term memory loss, difficulty processing information, sensory sensitivity, disturbed sleep pattern, and motor symptoms (e.g., twitching, poor coordination). Supplemental Figure 3 depicts the six symptoms that overlapped between both the ICC scale (rating day-of-scan severity 0- 10) and the medical history questionnaire (marking onset relative to acute COVID-19).

## 4. DISCUSSION

We conducted PET neuroimaging and blood analyses in individuals with ongoing diverse symptoms that began with acute COVID-19 infection. The majority of PASC participants’ acute COVID-19 illness did not require hospitalization. We found a significant increase in [^11^C]PBR28 signal, an indicator of neuroinflammation, across a wide area of brain regions such as the midcingulate cortex, corpus callosum, thalamus, basal ganglia/striatum, subfornical organ, anterior cingulate cortex, medial frontal gyrus, and precentral gyrus.

Of note, one study reported increased TSPO signal in moderately depressed PASC participants, 45% (9 of 20) of which had pre-COVID depression (Braga et al., 2023). In contrast, in the current study’s PASC cohort, the average depression score was mild, and no PASC participant reported a pre-COVID history of depression. Instead, they either denied depression being a problem for them (50%, 6 of 12) or reported that it was a new (∼42%, 5 of 12) or serious new (∼8%, 1 of 12) problem since COVID-19, and only 2 of 12 PASC participants were in the moderate depression severity range. Future studies should also include COVID-19-recovered individuals as a comparison group.

Importantly, in our study, we found that intensity of whole-brain PET signal showed significant positive correlations with blood measures related to vascular health (see Figure 2 and Table 3). This is indirect evidence that differences in PET signal across brain structures (Figure 1B) may partially reflect variance in vascular anatomy and perivascular immune penetration. To our knowledge, ours is the first study to provide direct evidence that processes related to neuroinflammation and vascular dysfunction are directly related in PASC.

As examples, fibrinogen and sL-selectin were two analytes from the vascular health multiplex panel that were significantly correlated with neuroinflammation-related PET signal. Elevated fibrinogen is associated with worse outcome in acute COVID-19 (Sui et al., 2021) and is not only associated with coagulation abnormalities in acute COVID-19 but also PASC (Pretorius et al., 2022; Sui et al., 2021). Related to neuroinflammation, fibrinogen persistently activates glia at the glymphatic (Mestre et al., 2017) perivascular spaces that surround neurovasculature (Davalos et al., 2012). Activated perivascular glia attract both glia from within the brain parenchyma and attract circulating immune factors to cross from neurovascular blood into brain. L-selectin (CD62L) is an adhesion molecule involved in attaching leukocytes (white blood cells) to vascular endothelium at sites of inflammation. Attraction of activated immune cells, inflammation of vascular endothelium, mobilization of glia, and activation of glia would each increase TSPO concentrations and therefore PET signal (as well as the symptoms associated with glial activation). The fact that these and other measures related to vascular dysfunction correlated with neuroinflammation provides evidence that the spatial pattern of PET signal in PASC vs. controls may reflect a vascular-related anatomical pattern that is likely to differ among patients. See Box 1 for further explanation of vascular analytes.

### Box 1

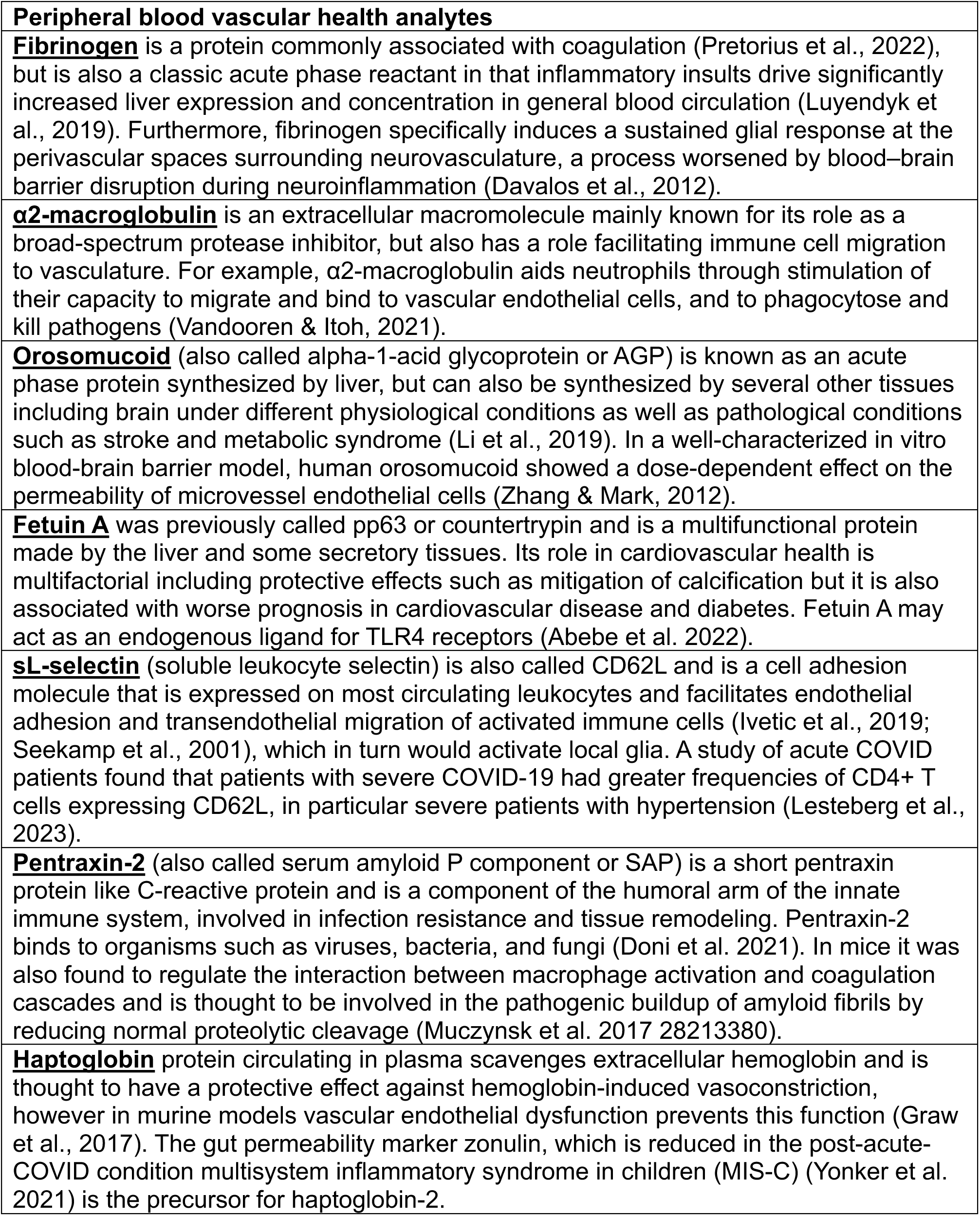

These vascular factors may penetrate into brain parenchyma via the perivascular spaces that line the neurovasculature and form the blood-brain barrier (Ineichen et al., 2022). During neuroinflammation, the blood-brain barrier becomes more permissive by opening up at the perivascular spaces, allowing circulating factors to affect the brain’s glia (Galea, 2021). We found increased PET uptake in some regions that tend to have dilated perivascular spaces. For example, we found significantly increased PET signal especially within left lentiform nucleus of the basal ganglia. The basal ganglia are a group of subcortical nuclei with many small, thin blood vessels that are common sites of unique perivascular anatomy and enlarged/dilated perivascular spaces (Mestre et al., 2017; Rudie et al., 2018), and apparently-subclinical neurovascular abnormalities such as cerebral microhemorrhages (Viswanathan & Chabriat, 2006), both of which have been reported in acute COVID-19 autopsy (Kantonen et al., 2020). In neuroradiology, dilated perivascular spaces on the lenticulostriate arteries projecting into the basal ganglia are relatively common and referred to as “Type 1.” Enlarged Type 1 perivascular spaces are sometimes seen in normal healthy aging but are associated with disease severity in some neurological conditions (Chan et al., 2021) and in one study were associated with sleep dysfunction in PASC (Del Brutto et al., 2022).

Within subcortical areas, some of the TSPO signal elevation pattern we measured also appears to be compatible with the location of the subfornical organ and the choroid plexus and ependymal glial cells at the roof of the third and floor of lateral ventricles, which are circumventricular organs (CVOs). CVOs are dense with ACE2 receptors (Ong et al., 2022), highly vascularized, and (except for the subcommissural organ) feature fenestrated blood vessels that lack a complete blood-brain barrier. This renders the glia near CVOs particularly vulnerable to being activated by blood-borne factors. We also detected elevated [^11^C]PBR28 PET signal across cingulate and corpus callosum, a pattern that follows the anterior cerebral artery pericallosal branch. More posterior regions of cingulate are particularly highly vascularized (Vogt, 2019). Unlike the CVOs but like the rest of the brain, these regions feature an intact blood brain barrier.

However, neurovascular endothelial cells are the blood-facing component of the blood-brain barrier and can be activated by circulating factors, in turn driving activation of glia (Kanda et al., 1995; Lécuyer et al., 2016; Pan et al., 2011). Furthermore, a reduced integrity of the blood-brain barrier, as observed during neuroinflammation (Galea, 2021), would likely contribute to this effect.

While our observational case-control study design cannot determine a causal relationship between neuroinflammation and vascular health, multiple potential non-mutually exclusive drivers (Proal & VanElzakker, 2021) of these effects are possible. Ongoing vascular-related problems and neuroinflammation may reflect lingering consequences from tissue injury during acute COVID-19. However most of our PASC participants did not report severe acute illness, and the acute COVID-19 illness that initiated their PASC was an average of 20.5 months prior to the scan visit. Under these circumstances, the ongoing objective neurological and vascular abnormalities may be more likely to reflect pre-COVID vulnerability factors and/or persistent stimulation by ongoing biological factors.

Previous studies have examined the contribution of other circulating factors in PASC. Vascular dysfunction may be consistent with the ongoing presence of fibrin-amyloid ’microclots’ (Pretorius et al., 2022) or with neutrophil hyperactivation (Boribong et al., 2022) that have been reported in individuals with PASC. These factors may represent continuous provocation by uncleared viral reservoirs at least in some PASC patients (Proal et al., 2023; Proal & VanElzakker, 2021). SARS-CoV-2 RNA or protein have been identified in PASC tissue after acute COVID-19, and multiple research groups have identified SARS-CoV-2 proteins including spike in blood in a subset of PASC individuals up to 16 months after initial infection (M. Peluso, 2023; Swank et al., 2023). These proteins may have leaked into general circulation from a tissue reservoir site via extracellular vesicle transport (Craddock et al., 2023; M. J. Peluso et al., 2022). Given that the spike protein drives coagulation cascades (Zheng et al., 2021) and a profound inflammatory response (e.g., Boribong et al., 2022), future PASC studies should measure these variables together in PASC cases versus COVID-recovered controls to better understand how they may be related to neuroinflammation and ongoing symptoms. Better understanding of these potential sources of neuroinflammation would be important for guiding related treatments and clinical trials.

### 4.2 LIMITATIONS

The current study compared 12 PASC patients to 43 controls, the majority of whom (34) were scanned pre-pandemic, the rest of whom tested negative for SARS-CoV-2 antibodies. Future studies should also recruit a specific well-defined PASC phenotype but increase the number of PASC participants, and should also compare PASC to COVID-19-recovered controls. While there was a higher proportion of female participants in the PASC group, we statistically controlled for sex in the primary analysis, and conducted a paired validation analysis in which PASC participants were directly matched for sex, genotype, and age (±5yrs). Furthermore, we conducted a confirmatory unpaired analysis comparing only female PASC cases to only female controls and found a very similar pattern (data not shown).

While the TSPO-targeting radioligand [^11^C]PBR28 is a widely used measure in studies of neuroinflammation, the specific function of TSPO is an area of ongoing study. In a non-mutually exclusive fashion, increased TSPO may reflect various activation states of glial cell types such as microglia and astrocytes (Paolicelli et al., 2022), density of glial cells (some of which are motile (Smolders et al., 2019)), peripheral immune cells penetrating into brain, or inflammatory activation of neurovascular endothelial cells (Guilarte et al., 2022). Therefore the exact biological driver of the observed increased PET signal in PASC remains to be fully elucidated.

Kinetic modeling using radiometabolite-corrected arterial input function, which is considered by many to be the gold-standard for TSPO signal quantification, was not included in the current study because the PASC participants did not undergo arterial line blood analysis during scanning and arterial line data were available only in a small subset of controls. While several previous studies have used arterial line blood-derived distribution volume (VT) ratio (DVR) to demonstrate the validity of SUVR as a semiquantitative ratio metric (e.g., (Alshelh et al., 2020)), future studies should include arterial line analyses in all participants or in a validation subset to further quantify the relationship between TSPO and measures of vascular health.

## Acknowledgments

Funding provided by PolyBio Research Foundation. Additional funding for control scans, Martinos Center infrastructure, and author effort provided by NIH grants 3R01DA047088-05S1, 1R21NS130283-01A1, 1S10RR023401, 1S10RR019307, and 1S10RR023043. Thanks to Brent Vogt PhD and Amy Proal PhD for helpful feedback on earlier versions of the manuscript, the Martinos Center PET and radiochemistry team, and especially the study participants.

## REFERENCES

Abebe, E. C., Tilahun Muche, Z., Behaile T/Mariam, A., Mengie Ayele, T., Mekonnen Agidew, M., Teshome Azezew, M., Abebe Zewde, E., Asmamaw Dejenie, T., & Asmamaw Mengstie, M. (2022). The structure, biosynthesis, and biological roles of fetuin-A: A review. Frontiers in Cell and Developmental Biology, 10, 945287. 10.3389/fcell.2022.945287

Ajčević, M., Iscra, K., Furlanis, G., Michelutti, M., Miladinović, A., Buoite Stella, A., Ukmar, M., Cova, M. A., Accardo, A., & Manganotti, P. (2023). Cerebral hypoperfusion in post-COVID- 19 cognitively impaired subjects revealed by arterial spin labeling MRI. Scientific Reports, 13(1), Article 1. 10.1038/s41598-023-32275-3

Albrecht, D. S., Forsberg, A., Sandstrom, A., Bergan, C., Kadetoff, D., Protsenko, E., Lampa, J., Lee, Y. C., Olgart Höglund, C., Catana, C., Cervenka, S., Akeju, O., Lekander, M., Cohen, G., Halldin, C., Taylor, N., Kim, M., Hooker, J. M., Edwards, R. R., … Loggia, M. L. (2019). Brain glial activation in fibromyalgia—A multi-site positron emission tomography investigation. Brain, Behavior, and Immunity, 75, 72–83. 10.1016/j.bbi.2018.09.018

Albrecht, D. S., Granziera, C., Hooker, J. M., & Loggia, M. L. (2016). In Vivo Imaging of Human Neuroinflammation. ACS Chemical Neuroscience, 7(4), 470–483. 10.1021/acschemneuro.6b00056

Alshelh, Z., Albrecht, D. S., Bergan, C., Akeju, O., Clauw, D. J., Conboy, L., Edwards, R. R., Kim, M., Lee, Y. C., Protsenko, E., Napadow, V., Sullivan, K., & Loggia, M. L. (2020). In-vivo imaging of neuroinflammation in Veterans with Gulf War Illness. Brain, Behavior, and Immunity, 87, 498–507. 10.1016/j.bbi.2020.01.020

Alshelh, Z., Brusaferri, L., Saha, A., Morrissey, E., Knight, P., Kim, M., Zhang, Y., Hooker, J. M., Albrecht, D., Torrado-Carvajal, A., Placzek, M. S., Akeju, O., Price, J., Edwards, R. R., Lee, J., Sclocco, R., Catana, C., Napadow, V., & Loggia, M. L. (2021). Neuroimmune signatures in chronic low back pain subtypes. Brain, 145(3), Article 3. 10.1093/brain/awab336

Bai, F., Tomasoni, D., Falcinella, C., Barbanotti, D., Castoldi, R., Mulè, G., Augello, M., Mondatore, D., Allegrini, M., Cona, A., Tesoro, D., Tagliaferri, G., Viganò, O., Suardi, E., Tincati, C., Beringheli, T., Varisco, B., Battistini, C. L., Piscopo, K., … Monforte, A. d’Arminio. (2022). Female gender is associated with long COVID syndrome: A prospective cohort study. Clinical Microbiology and Infection, 28(4), 611.e9-611.e16. 10.1016/j.cmi.2021.11.002

Barrientos, R. M., Higgins, E. A., Biedenkapp, J. C., Sprunger, D. B., Wright-Hardesty, K. J., Watkins, L. R., Rudy, J. W., & Maier, S. F. (2006). Peripheral infection and aging interact to impair hippocampal memory consolidation. Neurobiology of Aging, 27(5), 723–732. 10.1016/j.neurobiolaging.2005.03.010

Beck, A. T., Steer, R. A., Ball, R., & Ranieri, W. (1996). Comparison of Beck Depression Inventories-IA and -II in psychiatric outpatients. Journal of Personality Assessment, 67(3), 588–597. 10.1207/s15327752jpa6703_13

Berkman, J. M., Rosenthal, J. A., & Saadi, A. (2021). Carotid Physiology and Neck Restraints in Law Enforcement: Why Neurologists Need to Make Their Voices Heard. JAMA Neurology, 78(3), 267–268. 10.1001/jamaneurol.2020.4669

Boribong, B. P., LaSalle, T. J., Bartsch, Y. C., Ellett, F., Loiselle, M. E., Davis, J. P., Gonye, A. L. K., Sykes, D. B., Hajizadeh, S., Kreuzer, J., Pillai, S., Haas, W., Edlow, A. G., Fasano, A., Alter, G., Irimia, D., Sade-Feldman, M., & Yonker, L. M. (2022). Neutrophil profiles of pediatric COVID-19 and multisystem inflammatory syndrome in children. Cell Reports Medicine, 3(12), 100848. 10.1016/j.xcrm.2022.100848

Braga, J., Lepra, M., Kish, S. J., Rusjan, Pablo. M., Nasser, Z., Verhoeff, N., Vasdev, N., Bagby, M., Boileau, I., Husain, M. I., Kolla, N., Garcia, A., Chao, T., Mizrahi, R., Faiz, K., Vieira, E. L., & Meyer, J. H. (2023). Neuroinflammation After COVID-19 With Persistent Depressive and Cognitive Symptoms. JAMA Psychiatry, 80(8), 787–795. 10.1001/jamapsychiatry.2023.1321

Brusaferri, L., Alshelh, Z., Martins, D., Kim, M., Weerasekera, A., Housman, H., Morrissey, E. J., Knight, P. C., Castro-Blanco, K. A., Albrecht, D. S., Tseng, C.-E., Zürcher, N. R., Ratai, E.-M., Akeju, O., Makary, M. M., Catana, C., Mercaldo, N. D., Hadjikhani, N., Veronese, M., … Loggia, M. L. (2022). The pandemic brain: Neuroinflammation in non-infected individuals during the COVID-19 pandemic. Brain, Behavior, and Immunity, 102, 89–97. 10.1016/j.bbi.2022.02.018

Cabezas, R., Ávila, M., Gonzalez, J., El-Bachá, R. S., Báez, E., García-Segura, L. M., Jurado Coronel, J. C., Capani, F., Cardona-Gomez, G. P., & Barreto, G. E. (2014). Astrocytic modulation of blood brain barrier: Perspectives on Parkinson’s disease. Frontiers in Cellular Neuroscience, 8. https://www.frontiersin.org/articles/10.3389/fncel.2014.00211

Carruthers, B. M., van de Sande, M. I., De Meirleir, K. L., Klimas, N. G., Broderick, G., Mitchell, T., Staines, D., Powles, A. C. P., Speight, N., Vallings, R., Bateman, L., Baumgarten-Austrheim, B., Bell, D. S., Carlo-Stella, N., Chia, J., Darragh, A., Jo, D., Lewis, D., Light, A. R., … Stevens, S. (2011). Myalgic encephalomyelitis: International Consensus Criteria. Journal of Internal Medicine, 270(4), Article 4. 10.1111/j.1365-2796.2011.02428.x

Centers for Disease Control and Prevention. (2022). Long COVID or Post-COVID Conditions. https://www.cdc.gov/coronavirus/2019-ncov/long-term-effects/index.html

Chan, S. T., Mercaldo, N. D., Ravina, B., Hersch, S. M., & Rosas, H. D. (2021). Association of Dilated Perivascular Spaces and Disease Severity in Patients With Huntington Disease. Neurology, 96(6), Article 6. 10.1212/WNL.0000000000011121

Cleeland, C. S., & Ryan, K. M. (1994). Pain assessment: Global use of the Brief Pain Inventory. Annals of the Academy of Medicine, Singapore, 23(2), 129–138.

Craddock, V., Mahajan, A., Spikes, L., Krishnamachary, B., Ram, A. K., Kumar, A., Chen, L., Chalise, P., & Dhillon, N. K. (2023). Persistent circulation of soluble and extracellular vesicle-linked Spike protein in individuals with postacute sequelae of COVID-19. Journal of Medical Virology, 95(2), e28568. 10.1002/jmv.28568

Dantzer, R., & Kelley, K. W. (2007). Twenty Years of Research on Cytokine-Induced Sickness Behavior. Brain, Behavior, and Immunity, 21(2), 153–160. 10.1016/j.bbi.2006.09.006

Davalos, D., Kyu Ryu, J., Merlini, M., Baeten, K. M., Le Moan, N., Petersen, M. A., Deerinck, T. J., Smirnoff, D. S., Bedard, C., Hakozaki, H., Gonias Murray, S., Ling, J. B., Lassmann, H., Degen, J. L., Ellisman, M. H., & Akassoglou, K. (2012). Fibrinogen-induced perivascular microglial clustering is required for the development of axonal damage in neuroinflammation. Nature Communications, 3, 1227. 10.1038/ncomms2230

Davis, H. E., Assaf, G. S., McCorkell, L., Wei, H., Low, R. J., Re’em, Y., Redfield, S., Austin, J. P., & Akrami, A. (2021). Characterizing long COVID in an international cohort: 7 months of symptoms and their impact. EClinicalMedicine, 38, 101019. 10.1016/j.eclinm.2021.101019

Debruyne, J. C., Versijpt, J., Van Laere, K. J., De Vos, F., Keppens, J., Strijckmans, K., Achten, E., Slegers, G., Dierckx, R. A., Korf, J., & De Reuck, J. L. (2003). PET visualization of microglia in multiple sclerosis patients using [11C]PK11195. European Journal of Neurology, 10(3), Article 3. 10.1046/j.1468-1331.2003.00571.x

Del Brutto, O. H., Mera, R. M., Costa, A. F., Rumbea, D. A., Recalde, B. Y., & Castillo, P. R. (2022). Long coronavirus disease-related persistent poor sleep quality and progression of enlarged perivascular spaces. A longitudinal study. *Sleep*, zsac168. 10.1093/sleep/zsac168

Douaud, G., Lee, S., Alfaro-Almagro, F., Arthofer, C., Wang, C., McCarthy, P., Lange, F., Andersson, J. L. R., Griffanti, L., Duff, E., Jbabdi, S., Taschler, B., Keating, P., Winkler, A. M., Collins, R., Matthews, P. M., Allen, N., Miller, K. L., Nichols, T. E., & Smith, S. M. (2022). SARS-CoV-2 is associated with changes in brain structure in UK Biobank. Nature, 604(7907), Article 7907. 10.1038/s41586-022-04569-5

Galea, I. (2021). The blood–brain barrier in systemic infection and inflammation. Cellular & Molecular Immunology, 18(11), Article 11. 10.1038/s41423-021-00757-x

Graw, J. A., Yu, B., Rezoagli, E., Warren, H. S., Buys, E. S., Bloch, D. B., & Zapol, W. M. (2017). Endothelial dysfunction inhibits the ability of haptoglobin to prevent hemoglobin-induced hypertension. American Journal of Physiology - Heart and Circulatory Physiology, 312(6), H1120–H1127. 10.1152/ajpheart.00851.2016

Guilarte, T. R., Rodichkin, A. N., McGlothan, J. L., De La Rocha, A. M. A., & Azzam, D. J. (2022). Imaging neuroinflammation with TSPO: A new perspective on the cellular sources and subcellular localization. Pharmacology & Therapeutics, 234, 108048. 10.1016/j.pharmthera.2021.108048

Ineichen, B. V., Okar, S. V., Proulx, S. T., Engelhardt, B., Lassmann, H., & Reich, D. S. (2022). Perivascular spaces and their role in neuroinflammation. Neuron, 110(21), 3566–3581. 10.1016/j.neuron.2022.10.024

Ivetic, A., Hoskins Green, H. L., & Hart, S. J. (2019). L-selectin: A Major Regulator of Leukocyte Adhesion, Migration and Signaling. Frontiers in Immunology, 10, 1068. 10.3389/fimmu.2019.01068

Izquierdo-Garcia, D., Hansen, A. E., Förster, S., Benoit, D., Schachoff, S., Fürst, S., Chen, K. T., Chonde, D. B., & Catana, C. (2014). An SPM8-based Approach for Attenuation Correction Combining Segmentation and Non-rigid Template Formation: Application to Simultaneous PET/MR Brain Imaging. Journal of Nuclear Medicine : Official Publication, Society of Nuclear Medicine, 55(11), Article 11. 10.2967/jnumed.113.136341

Kalk, N. j., Owen, D. r., Tyacke, R. j., Reynolds, R., Rabiner, E. a., Lingford-hughes, A. r., & Parker, C. a. (2013). Are prescribed benzodiazepines likely to affect the availability of the 18 kDa translocator protein (TSPO) in PET studies? Synapse, 67(12), 909–912. 10.1002/syn.21681

Kanda, T., Yamawaki, M., Ariga, T., & Yu, R. K. (1995). Interleukin 1 beta up-regulates the expression of sulfoglucuronosyl paragloboside, a ligand for L-selectin, in brain microvascular endothelial cells. Proceedings of the National Academy of Sciences of the United States of America, 92(17), 7897–7901.

Kantonen, J., Mahzabin, S., Mäyränpää, M. I., Tynninen, O., Paetau, A., Andersson, N., Sajantila, A., Vapalahti, O., Carpén, O., Kekäläinen, E., Kantele, A., & Myllykangas, L. (2020). Neuropathologic features of four autopsied COVID-19 patients. Brain Pathology, 30(6), 1012–1016. 10.1111/bpa.12889

Klein, J., Wood, J., Jaycox, J., Lu, P., Dhodapkar, R. M., Gehlhausen, J. R., Tabachnikova, A., Tabacof, L., Malik, A. A., Kamath, K., Greene, K., Monteiro, V. S., Peña-Hernandez, M., Mao, T., Bhattacharjee, B., Takahashi, T., Lucas, C., Silva, J., Mccarthy, D., … Iwasaki, A. (2022). (Pre-print) Distinguishing features of Long COVID identified through immune profiling. medRxiv. 10.1101/2022.08.09.22278592

Lécuyer, M.-A., Kebir, H., & Prat, A. (2016). Glial influences on BBB functions and molecular players in immune cell trafficking. Biochimica et Biophysica Acta (BBA) - Molecular Basis of Disease, 1862(3), 472–482. 10.1016/j.bbadis.2015.10.004

Lesteberg, K. E., Araya, P., Waugh, K. A., Chauhan, L., Espinosa, J. M., & Beckham, J. D. (2023). Severely ill and high-risk COVID-19 patients exhibit increased peripheral circulation of CD62L+ and perforin+ T cells. Frontiers in Immunology, 14. https://www.frontiersin.org/articles/10.3389/fimmu.2023.1113932

Lindgren, N., Tuisku, J., Vuoksimaa, E., Helin, S., Karrasch, M., Marjamäki, P., Kaprio, J., & Rinne, J. O. (2020). Association of neuroinflammation with episodic memory: A [11C]PBR28 PET study in cognitively discordant twin pairs. Brain Communications, 2(1), fcaa024. 10.1093/braincomms/fcaa024

Líška, D., Liptaková, E., Babičová, A., Batalik, L., Baňárová, P. S., & Dobrodenková, S. (2022). What is the quality of life in patients with long COVID compared to a healthy control group? Frontiers in Public Health, 10. https://www.frontiersin.org/articles/10.3389/fpubh.2022.975992

Loggia, M. L., Chonde, D. B., Akeju, O., Arabasz, G., Catana, C., Edwards, R. R., Hill, E., Hsu, S., Izquierdo-Garcia, D., Ji, R.-R., Riley, M., Wasan, A. D., Zürcher, N. R., Albrecht, D. S., Vangel, M. G., Rosen, B. R., Napadow, V., & Hooker, J. M. (2015). Evidence for brain glial activation in chronic pain patients. Brain, 138(3), 604–615. 10.1093/brain/awu377

Luyendyk, J. P., Schoenecker, J. G., & Flick, M. J. (2019). The multifaceted role of fibrinogen in tissue injury and inflammation. Blood, 133(6), 511–520. 10.1182/blood-2018-07-818211

Ma, Y., Deng, J., Liu, Q., Du, M., Liu, M., & Liu, J. (2023). Long-Term Consequences of Asymptomatic SARS-CoV-2 Infection: A Systematic Review and Meta-Analysis. International Journal of Environmental Research and Public Health, 20(2), 1613. 10.3390/ijerph20021613

Mestre, H., Kostrikov, S., Mehta, R. I., & Nedergaard, M. (2017). Perivascular Spaces, Glymphatic Dysfunction, and Small Vessel Disease. Clinical Science (London, England : 1979), 131(17), 2257–2274. 10.1042/CS20160381

National Center for Health Statistics. (2023). U.S. Census Bureau, Household Pulse Survey, 2022- 2023. Long COVID. https://www.cdc.gov/nchs/covid19/pulse/long-covid.htm

Ong, W.-Y., Satish, R. L., & Herr, D. R. (2022). ACE2, Circumventricular Organs and the Hypothalamus, and COVID-19. Neuromolecular Medicine, 24(4), 363–373. 10.1007/s12017-022-08706-1

Owen, D. R., Guo, Q., Rabiner, E. A., & Gunn, R. N. (2015). The impact of the rs6971 polymorphism in TSPO for quantification and study design. Clinical and Translational Imaging, 3(6), 417–422. 10.1007/s40336-015-0141-z

Owen, D. R., Yeo, A. J., Gunn, R. N., Song, K., Wadsworth, G., Lewis, A., Rhodes, C., Pulford, D. J., Bennacef, I., Parker, C. A., StJean, P. L., Cardon, L. R., Mooser, V. E., Matthews, P. M., Rabiner, E. A., & Rubio, J. P. (2012). An 18-kDa Translocator Protein (TSPO) polymorphism explains differences in binding affinity of the PET radioligand PBR28. Journal of Cerebral Blood Flow & Metabolism, 32(1), 1–5. 10.1038/jcbfm.2011.147

Pan, W., Stone, K. P., Hsuchou, H., Manda, V. K., Zhang, Y., & Kastin, A. J. (2011). Cytokine Signaling Modulates Blood-Brain Barrier Function. Current Pharmaceutical Design, 17(33), 3729–3740.

Pannell, M., Economopoulos, V., Wilson, T. C., Kersemans, V., Isenegger, P. G., Larkin, J. R., Smart, S., Gilchrist, S., Gouverneur, V., & Sibson, N. R. (2020). Imaging of translocator protein upregulation is selective for pro-inflammatory polarized astrocytes and microglia. Glia, 68(2), 280–297. 10.1002/glia.23716

Paolicelli, R. C., Sierra, A., Stevens, B., Tremblay, M.-E., Aguzzi, A., Ajami, B., Amit, I., Audinat, E., Bechmann, I., Bennett, M., Bennett, F., Bessis, A., Biber, K., Bilbo, S., Blurton-Jones, M., Boddeke, E., Brites, D., Brône, B., Brown, G. C., … Wyss-Coray, T. (2022). Microglia states and nomenclature: A field at its crossroads. Neuron, 110(21), 3458–3483. 10.1016/j.neuron.2022.10.020

Patel, M. A., Knauer, M. J., Nicholson, M., Daley, M., Van Nynatten, L. R., Martin, C., Patterson, E. K., Cepinskas, G., Seney, S. L., Dobretzberger, V., Miholits, M., Webb, B., & Fraser, D. D. (2022). Elevated vascular transformation blood biomarkers in Long-COVID indicate angiogenesis as a key pathophysiological mechanism. Molecular Medicine, 28(1), 122. 10.1186/s10020-022-00548-8

Peluso, M. (2023). PLASMA-BASED ANTIGEN PERSISTENCE IN THE POST-ACUTE PHASE OF SARS-CoV-2 INFECTION - CROI Conference. https://www.croiconference.org/abstract/plasma-based-antigen-persistence-in-the-post-acute-phase-of-sars-cov-2-infection/

Peluso, M. J., Deeks, S. G., Mustapic, M., Kapogiannis, D., Henrich, T. J., Lu, S., Goldberg, S. A., Hoh, R., Chen, J. Y., Martinez, E. O., Kelly, J. D., Martin, J. N., & Goetzl, E. J. (2022). SARS-CoV-2 and Mitochondrial Proteins in Neural-Derived Exosomes of COVID-19. Annals of Neurology, 91(6), 772–781. 10.1002/ana.26350

Pretorius, E., Venter, C., Laubscher, G. J., Kotze, M. J., Oladejo, S. O., Watson, L. R., Rajaratnam, K., Watson, B. W., & Kell, D. B. (2022). Prevalence of symptoms, comorbidities, fibrin amyloid microclots and platelet pathology in individuals with Long COVID/Post-Acute Sequelae of COVID-19 (PASC). Cardiovascular Diabetology, 21(1), 148. 10.1186/s12933-022-01579-5

Proal, A. D., & VanElzakker, M. B. (2021). Long COVID or Post-acute Sequelae of COVID-19 (PASC): An Overview of Biological Factors That May Contribute to Persistent Symptoms. Frontiers in Microbiology, 12, 698169. 10.3389/fmicb.2021.698169

Proal, A. D., VanElzakker, M. B., Aleman, S., Bach, K., Boribong, B. P., Buggert, M., Cherry, S., Chertow, D. S., Davies, H. E., Dupont, C. L., Deeks, S. G., Eimer, W., Ely, E. W., Fasano, A., Freire, M., Geng, L. N., Griffin, D. E., Henrich, T. J., Iwasaki, A., … Wherry, E. J. (2023). SARS-CoV-2 reservoir in post-acute sequelae of COVID-19 (PASC). Nature Immunology, 1–12. 10.1038/s41590-023-01601-2

Rudie, J. D., Rauschecker, A. M., Nabavizadeh, S. A., & Mohan, S. (2018). Neuroimaging of Dilated Perivascular Spaces: From Benign and Pathologic Causes to Mimics. Journal of Neuroimaging : Official Journal of the American Society of Neuroimaging, 28(2), 139– 149. 10.1111/jon.12493

Seekamp, A., van Griensven, M., Hildebrandt, F., Brauer, N., Jochum, M., & Martin, M. (2001). The effect of trauma on neutrophil L-selectin expression and sL-selectin serum levels. *Shock (Augusta*, Ga.), 15(4), 254–260. 10.1097/00024382-200115040-00002

Smolders, S. M.-T., Kessels, S., Vangansewinkel, T., Rigo, J.-M., Legendre, P., & Brône, B. (2019). Microglia: Brain cells on the move. Progress in Neurobiology, 178, 101612. 10.1016/j.pneurobio.2019.04.001

Sui, J., Noubouossie, D. F., Gandotra, S., & Cao, L. (2021). Elevated Plasma Fibrinogen Is Associated With Excessive Inflammation and Disease Severity in COVID-19 Patients. Frontiers in Cellular and Infection Microbiology, 11. https://www.frontiersin.org/articles/10.3389/fcimb.2021.734005

Swank, Z., Senussi, Y., Manickas-Hill, Z., Yu, X. G., Li, J. Z., Alter, G., & Walt, D. R. (2023). Persistent Circulating Severe Acute Respiratory Syndrome Coronavirus 2 Spike Is Associated With Post-acute Coronavirus Disease 2019 Sequelae. Clinical Infectious Diseases: An Official Publication of the Infectious Diseases Society of America, 76(3), e487–e490. 10.1093/cid/ciac722

Tayal, S., Ali, A., Kumar, V., Jha, A. K., & Gandhi, A. (2021). Importance of Understanding and Analyzing Daily Quality Assurance Test of Positron Emission Tomography/Computed Tomography Equipment in Minimizing the Downtime of Equipment in Remote Places. *Indian Journal of Nuclear Medicine : IJNM : The Official Journal of the Society of Nuclear Medicine*, India, 36(2), 179–182. 10.4103/ijnm.IJNM_196_20

Tzourio-Mazoyer, N., Landeau, B., Papathanassiou, D., Crivello, F., Etard, O., Delcroix, N., Mazoyer, B., & Joliot, M. (2002). Automated anatomical labeling of activations in SPM using a macroscopic anatomical parcellation of the MNI MRI single-subject brain. NeuroImage, 15(1), 273–289. 10.1006/nimg.2001.0978

Vandooren, J., & Itoh, Y. (2021). Alpha-2-Macroglobulin in Inflammation, Immunity and Infections. Frontiers in Immunology, 12. https://www.frontiersin.org/articles/10.3389/fimmu.2021.803244

VanElzakker, M. B., Brumfield, S. A., & Lara Mejia, P. S. (2019). Neuroinflammation and Cytokines in Myalgic Encephalomyelitis/Chronic Fatigue Syndrome (ME/CFS): A Critical Review of Research Methods. Frontiers in Neurology, 9, 1033. 10.3389/fneur.2018.01033

Varga, Z., Flammer, A. J., Steiger, P., Haberecker, M., Andermatt, R., Zinkernagel, A. S., Mehra, M. R., Schuepbach, R. A., Ruschitzka, F., & Moch, H. (2020). Endothelial cell infection and endotheliitis in COVID-19. *Lancet (London*, England), 395(10234), 1417–1418. 10.1016/S0140-6736(20)30937-5

Viswanathan, A., & Chabriat, H. (2006). Cerebral Microhemorrhage. Stroke, 37(2), 550–555. 10.1161/01.STR.0000199847.96188.12

Vogt, B. A. (2019). Chapter 1 - The cingulate cortex in neurologic diseases: History, Structure, Overview. In B. A. Vogt (Ed.), Handbook of Clinical Neurology (Vol. 166, pp. 3–21). Elsevier. 10.1016/B978-0-444-64196-0.00001-7

Winkler, A. M., Ridgway, G. R., Webster, M. A., Smith, S. M., & Nichols, T. E. (2014). Permutation inference for the general linear model. Neuroimage, 92(100), 381–397. 10.1016/j.neuroimage.2014.01.060

Woolrich, M. W., Behrens, T. E. J., Beckmann, C. F., Jenkinson, M., & Smith, S. M. (2004). Multilevel linear modelling for FMRI group analysis using Bayesian inference. NeuroImage, 21(4), 1732–1747. 10.1016/j.neuroimage.2003.12.023

World Health Organization. (2021). A clinical case definition of post COVID-19 condition by a Delphi consensus, 6 October 2021. https://www.who.int/publications/i/item/WHO-2019-nCoV-Post_COVID-19_condition-Clinical_case_definition-2021.1

Xie, Y., Xu, E., Bowe, B., & Al-Aly, Z. (2022). Long-term cardiovascular outcomes of COVID-19. Nature Medicine, 28(3), Article 3. 10.1038/s41591-022-01689-3

Xu, E., Xie, Y., & Al-Aly, Z. (2022). Long-term neurologic outcomes of COVID-19. Nature Medicine, 28(11), Article 11. 10.1038/s41591-022-02001-z

Yonker, L. M., Gilboa, T., Ogata, A. F., Senussi, Y., Lazarovits, R., Boribong, B. P., Bartsch, Y. C., Loiselle, M., Rivas, M. N., Porritt, R. A., Lima, R., Davis, J. P., Farkas, E. J., Burns, M. D., Young, N., Mahajan, V. S., Hajizadeh, S., Lopez, X. I. H., Kreuzer, J., … Fasano, A. (2021). Multisystem inflammatory syndrome in children is driven by zonulin-dependent loss of gut mucosal barrier. The Journal of Clinical Investigation, 131(14), e149633. 10.1172/JCI149633

Zeng, N., Zhao, Y.-M., Yan, W., Li, C., Lu, Q.-D., Liu, L., Ni, S.-Y., Mei, H., Yuan, K., Shi, L., Li, P., Fan, T.-T., Yuan, J.-L., Vitiello, M. V., Kosten, T., Kondratiuk, A. L., Sun, H.-Q., Tang, X.-D., Liu, M.- Y., … Lu, L. (2023). A systematic review and meta-analysis of long term physical and mental sequelae of COVID-19 pandemic: Call for research priority and action. Molecular Psychiatry, 28(1), 423–433. 10.1038/s41380-022-01614-7

Zheng, Y., Zhao, J., Li, J., Guo, Z., Sheng, J., Ye, X., Jin, G., Wang, C., Chai, W., Yan, J., Liu, D., & Liang, X. (2021). SARS-CoV-2 spike protein causes blood coagulation and thrombosis by competitive binding to heparan sulfate. International Journal of Biological Macromolecules, 193, 1124–1129. 10.1016/j.ijbiomac.2021.10.112

