## Supplemental data for "Neuroinflammation in post-acute sequelae of COVID-19 (PASC) as assessed by [^11^C]PBR28 PET correlates with vascular disease measures"

### SUPPLEMENTAL FIGURES

Supplemental Figure 1

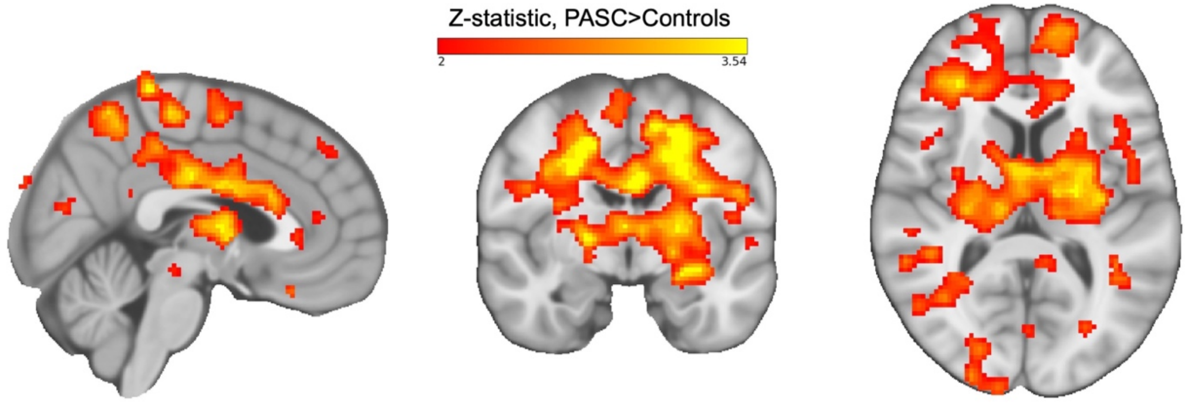


Supplemental Figure 1 legend:

To validate our primary results, we performed a second analysis using a paired approach in which each PASC participant was matched to a control participant for genotype, sex, and age ±5 years. Because no control participant fit these criteria for one PASC participant, they were left out of the paired validation analysis. Genotype and sex were directly matched; neither age nor cerebellum SUV differed between cases and controls ($p>0.05$). Example slices through sagittal (x=3), coronal (y=-7), and axial (z=8) sections of the 11 PASC > 11 control group-level unpaired validation comparison, showing the pattern of increased [$^{11}$C]PBR28 signal. Color bar: threshold min. Z score of 2 and max. 3.54. (Figure is shown in neurological convention)

Supplemental Figure 2

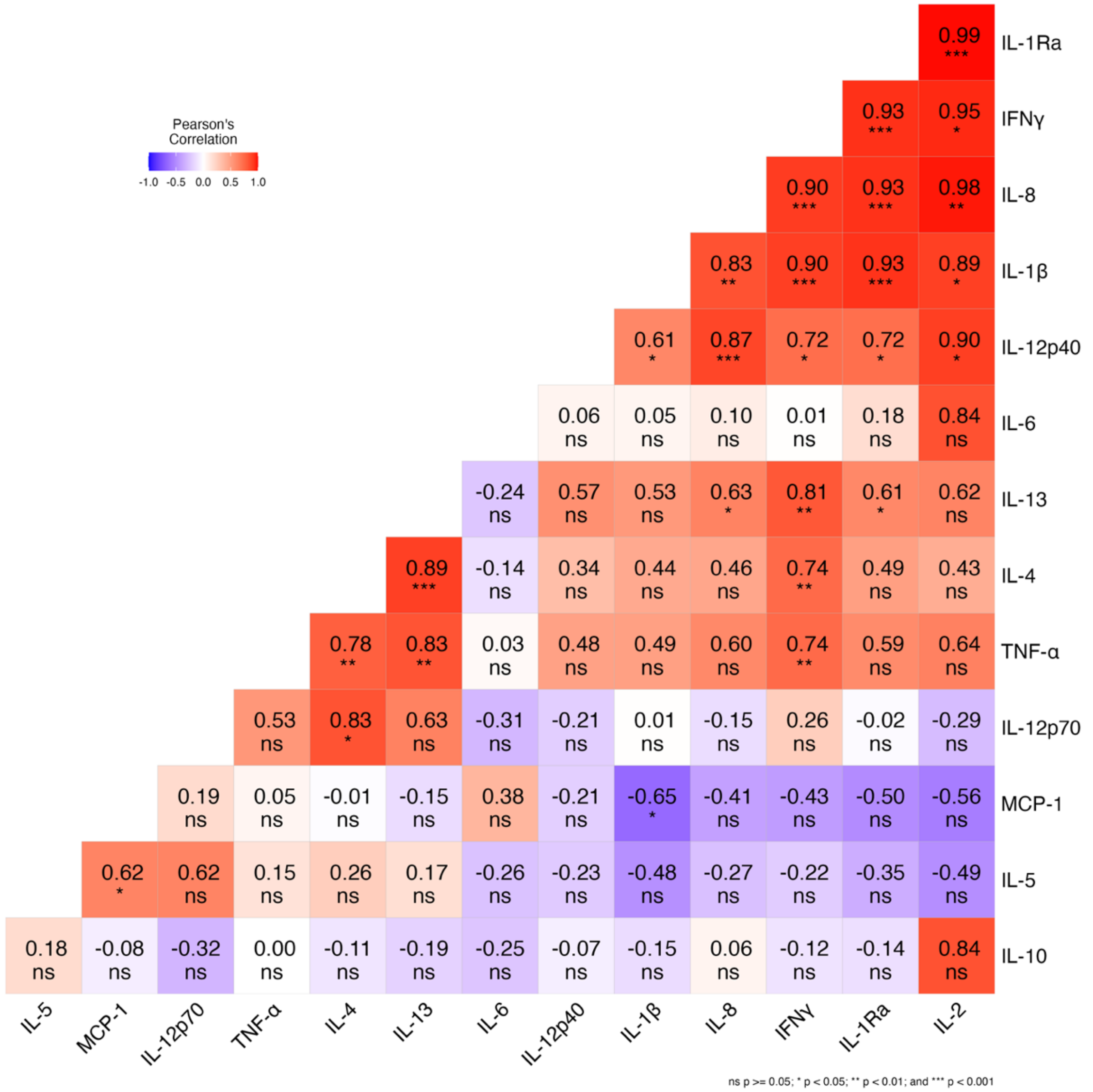


Supplemental Figure 2 legend:

Correlation matrix showing relationships among cytokines from a 15 cytokine multiplex (Millipore HCYTA-60K Luminex magnetic bead panel performed by Eve Technologies, Calgary Canada), from the 11 PASC participants that provided platelet-poor plasma samples immediately before PET neuroimaging.

Analytes included: GM-CSF, IFNγ, IL-1β, IL-1RA, IL-2, IL-4, IL-5, IL-6, IL-8, IL-10, IL-12(p40), IL-12(p70), IL-13, MCP-1, TNFα. * = $p<0.05$; ** = $p<0.01$; *** = $p<0.001$

Supplemental Figure 3

| PASC participant | 1 | 2 | 3 | 4 | 5 | 6 | 7 | 8 | 9 | 10 | 11 | 12 |
|---|---|---|---|---|---|---|---|---|---|---|---|---|
| Headache | 9 | 8 | 0 | 2 | 0 | 7 | 10 | 6 | 0 | 9 | 6 | 2 |
| Unrested | 10 | 8 | 4 | 6 | 1 | 0 | 10 | 9 | 7 | 5 | 10 | 7 |
| Muscle / Joint } Pain | 10 | 3 | 0 | 2 | 6 | 6 | 10 | 6 | 5 | 1 | 3 | 2 |
| Short Term Memory | 7 | 6 | 6 | 4 | 4 | 0 | 10 | 0 | 2 | 6 | 2 | 7 |
| Processing Information | 9 | 6 | 6 | 6 | 4 | 0 | 10 | 3 | 3 | 6 | 3 | 5 |

Supplemental Figure 3 legend:

Each of the 12 PASC participants rated symptoms from the ICC “Neurological impairments” cluster on a 1-10 severity scale and marked their onset. For Pain, the ICC asked about "Significant pain" but the history questionnaire distinguished between muscle and joint pain.

Light green = Not reported as a problem

Orange = “This was a problem for me before COVID"

Pink = "This is a problem for me since I had COVID"

Red = "This is a serious problem for me since I had COVID"

Supplemental Table 1

| **Affective items** | PASC mean (SD) | Control mean (SD) |
|---|---|---|
| Sadness | 0.58 (0.67) | 0.095 (0.48) |
| Pessimism | 0.67 (0.49) | 0.024 (0.15) |
| Sense of failure | 0.00 (0.00) | 0.024 (0.15) |
| Dissatisfaction | 0.67 (0.65) | 0.095 (0.30) |
| Guilt | 0.17 (0.39) | 0.023 (0.15) |
| Feeling of punishment | 0.00 (0.00) | 0.048 (0.22) |
| Disappointment in self | 0.17 (0.39) | 0.024 (0.16) |
| Self-criticism | 0.58 (0.51) | 0.048 (0.22) |
| Suicidal thoughts | 0.083 (0.29) | 0.00 (0.00) |
| Crying episodes | 0.42 (0.90) | 0.048 (0.22) |
| Irritability | 0.42 (0.51) | 0.24 (0.69) |
| Social withdrawal | 0.50 (0.52) | 0.048 (0.22) |
| Indecisiveness | 1.00 (0.74) | 0.00 (0.00) |
| **Average affective items** | **0.40 (0.23)** | **0.055 (0.10)** |
| **Somatic Items** | | |
| Change of body image | 0.42 (0.51) | 0.024 (0.15) |
| Ability to work | 1.17 (0.94) | 0.073 (0.26) |
| Sleep difficulties | 0.75 (0.45) | 0.17 (0.44) |
| Fatigability | 1.42 (0.79) | 0.073 (0.26) |

| | | |
|---|---|---|
| Loss of Appetite | 0.58 (0.79) | 0.024 (0.15) |
| Weight loss | 0.083 (0.29) | 0.048 (0.22) |
| Health anxiety | 1.08 (0.67) | 0.00 (0.00) |
| Loss of libido | 0.83 (1.03) | 0.071 (0.26) |
| **Average somatic items** | **0.79 (0.39)** | **0.060 (0.13)** |
| **Total score averages** | **11.58 (1.74)** | **1.14 (5.58)** |

Supplemental Table 1 legend:

PASC and Control participant means for BDI items within the affective and somatic subscales of the BDI. A mixed-model ANOVA revealed a significant interaction driven by high somatic item means within the PASC group.
